# Neural basis for anxiety and anxiety-related physiological responses during a driving situation: An fMRI study

**DOI:** 10.1101/2021.11.30.470539

**Authors:** Takafumi Sasaoka, Tokiko Harada, Daichi Sato, Nanae Michida, Hironobu Yonezawa, Masatoshi Takayama, Takahide Nouzawa, Shigeto Yamawaki

**Author notes:** **Corresponding author:** Takafumi Sasaoka, PhD, Center for Brain, Mind, and KANSEI Sciences Research, Hiroshima University, 1-2-3 Kasumi, Minami-ku, Hiroshima, 734-8551, Japan.

## Abstract

While the exteroceptive and interoceptive prediction of a negative event increases a person’s anxiety in daily life situations, the relationship between the brain mechanism of anxiety and the anxiety-related autonomic response has not been fully understood. In this fMRI study, we examined the neural basis of anxiety and anxiety-related autonomic responses in a daily driving situation. Participants viewed a driving video clip in the first-person perspective. During the video clip, participants were presented with a cue to indicate whether a subsequent crash could occur (attention condition) or not (safe condition). Enhanced activities in the anterior insula, bed nucleus of the stria terminalis, thalamus, and periaqueductal gray, and higher sympathetic nerve responses (pupil dilation and peripheral arterial stiffness) were triggered by the attention condition but not with the safe condition. Autonomic response-related functional connectivity was detected in the visual cortex, cerebellum, brainstem, and MCC/PCC with the right anterior insula and its adjacent regions as seed regions. Thus, the right anterior insula and adjacent regions, in collaboration with other regions play a role in eliciting anxiety based on the prediction of negative events, by mediating anxiety-related autonomic responses according to interoceptive information.

## Introduction

It has been emphasized that the fundamental function of the brain, as a “prediction machine,” is prediction and prediction error processing (e.g., Clark 2013). The predictive coding theory (Friston 2005; Rao and Ballard 1999) has been proposed as a computational framework for brain prediction and prediction error processing. According to this theory, the brain works on the computational principle to minimize the prediction error (difference between the prediction of the cause of the sensory input and the actual sensory input). Thus, the cause of the sensory input is inferred: to minimize prediction errors, predictions are updated and actions are performed in order to change the sensory input and to confirm the predictions (active inference). Based on this theory, perception could be regarded as an inference of the cause of external sensation (i.e., exteroception) of the outer world by prediction and prediction-error processing of exteroception (Rao and Ballard 1999). Likewise, subjective feelings could be regarded as an inference of the cause of the bodily sensation (i.e., interoception) by prediction and prediction-error processing of interoception (e.g., Seth 2013, Barrett and Simmons 2015; Barrett et al. 2016; Barrett 2017).

Anxiety, a typical example of brain prediction processing, can be defined by anticipatory affective, cognitive, and behavioral changes in response to uncertainty regarding potential threats (Grupe and Nitshke 2013). When the brain predicts a threatening event that can disturb homeostasis, negative emotions are elicited and attention is directed to the incoming information; thus, actions are undertaken to avoid the event. This corresponds to a state of anxiety. The interoception processing model based on the predictive coding theory (Seth 2013) assumes that the prediction of upcoming threatening events results in updating the desired state of the organism, accompanied by autonomic reflexes. Based on this assumption, we can predict that the elicitation of anxiety will trigger a corresponding autonomic response. Understanding the neural basis for eliciting anxiety and accompanying autonomic responses in a daily situation will contribute to understanding the mechanisms of anxiety and anxiety disorders, as well as to implementing a system to appropriately reduce anxiety by reading out anxiety-related autonomic responses.

Previous neuroimaging studies on anxiety disorder and phobia have suggested that the anterior insula and amygdala play a key role in these conditions. A meta-analysis of neuroimaging studies on anxiety disorder (Etkin and Wager 2007) showed that hyperactivation in the amygdala and insula was more frequently related to social anxiety disorder and specific phobia than to posttraumatic stress disorder. In line with this, Baur et al. (2013) demonstrated that state and trait anxiety were associated with the functional and structural connectivity between the anterior insula and the amygdala. Specific phobias, which are prevalent anxiety disorders, are associated with activation in the brain network (“fear network”; Sehlmeyer et al. 2009), including the medial prefrontal cortex, anterior cingulate cortex (ACC), insula, and amygdala. Indeed, Schweckendiek et al. (2011) investigated fear learning in arachnophobia using a fear conditioning paradigm. Their study showed that arachnophobia was linked to enhanced activation in the fear network including the medial prefrontal cortex, ACC, insula, thalamus, and amygdala. Moreover, they showed increased amygdala activation in response to the phobia-related conditioned stimulus (CS), but not in response to the non-phobia-related CS. A meta-analysis of fear conditioning neuroimaging studies (Fullana et al. 2016) showed that the fear CS consistently activated the “central autonomic-interoceptive network,” which includes the anterior insula, dorsal ACC (dACC), and the subcortical viscero-sensory regions such as the midbrain (periaqueductal gray [PAG] and parabrachial nucleus), ventral thalamus, hypothalamus, and the pontomedullary junction. This activation reflects the involvement of the conscious experience of fear and anxiety and the autonomic responses to threat. The amygdala involvement in the fear conditioning was not apparent in the meta-analysis study. On the other hand, it has been reported that the extended amygdala, including the bed nucleus of the stria terminalis (BNST), plays a crucial role in anxiety. Indeed, Somerville et al. (2013) conducted an fMRI study showing that the BNST is associated with the sustained response anticipating the aversive stimuli, whereas the amygdala is involved with the transient response to such stimuli. Therefore, it is possible that the amygdala and the extended amygdala have different roles in anxiety elicitation: the amygdala activity is related to fear itself, whereas the BNST is more related to the anticipation of fearful events. In contrast, the default-mode network (DMN), including the ventral medial prefrontal cortex and posterior cingulate cortex (PCC), was deactivated when CS was presented relative to non-CS (“safe-signal”) (Fullana et al. 2016). This could yield various interpretations, such as fear-response inhibition, encoding episodic memory traces of the conditioned-unconditioned stimuli (CS/US) association, and relief related to omission of the US. However, the role of the safe-signal-related brain network remains unclear.

In the theory of anxiety disorders, some researchers have emphasized the deficit of the interoceptive processing with the ability to mitigate anxiety. Paulus and Stein (2006) proposed that individuals who are prone to anxiety show an altered interoceptive prediction in the anterior insula. For the brain, excessive interoceptive prediction could need additional resources to reduce the prediction error, resulting in increased anxiety in anticipation of aversive events. Interoceptive information is often first processed in the brainstem, such as the medial nucleus of the solitary tract and the parabrachial nucleus, and relayed to the insula by the thalamus (Chen et al. 2021). The anterior insula has been suggested to be involved in the integration of interoceptive and exteroceptive representations, as well as in their updating based on the incoming information (Craig 2009). Based on these observations, we can speculate that perception of a cue associated with threatening events triggers a prediction of negative body states generated by the anterior insula, and that accompanying interoceptive information is processed by the subcortical regions, such as the thalamus and brainstem.

It is well known that autonomic responses, such as increasing heart rate (e.g., Deane 1961), skin conductance (e.g., Epstein and Roupenian 1970), and pupil dilation (e.g., Bitsios et al. 2004) are observed when people experience anxiety. However, there are few studies examining the brain mechanisms involved in the autonomic responses related to anxiety. For instance, Wager et al. (2009a, 2009b) reported that the dorsal pregenual cingulate region was active in relation to the increase in the heart rate while facing a socially threatening situation in which participants were waiting to deliver a speech to others. However, many other indices reflect sympathetic nerve activity rather than the heart rate. Peripheral arterial stiffness (β_art_) has been proposed (Matsubara et al. 2018; Tsuji et al. 2021) as an index reflecting sympathetic activity. β_art_ can be estimated using electrocardiography, continuous sphygmomanometry, and photoplethysmography (PPG). Tsuji et al. (2021) demonstrated that β_art_ correlated with subjective pain ratings and brain activity in regions including the salience network (e.g., dACC), suggesting that β_art_ is a potential candidate for a sympathetic nerve index reflecting anxiety. Cardiovascular autonomic signals are often used as autonomic indices during driving, especially as a measure of mental workload (Paxion et al. 2014). For example, heart rate and a specific component of heart rate variability (0.10 Hz) reflect the mental workload during driving (e.g., Brookhuis and De Waard 1991; Liu and Lee 2006).

In the current study, we examined the brain mechanisms involved in eliciting anxiety and related to induced autonomic responses by using a situation of driving, one of the possible situations that would frequently cause anxiety in daily life. We performed simultaneous measurements via functional magnetic resonance imaging (fMRI) and other physiological indices, including electrocardiography (ECG), PPG, blood pressure monitoring, and pupillometry. In particular, we focused on pupil dilation and β_art_ as putative indications of sympathetic nerve activity. We hypothesized that the anterior insula and subcortical regions involved in threat and interoceptive processing (BNST, thalamus, and midbrain), are active in anticipation of the crash. We anticipated that if the amygdala was involved in the actual threatening event, its activity would increase in response to the crash, and that enhanced BNST activity would be detected in anticipation of the crash. Moreover, we predicted significant physiological responses indicating sympathetic nerve activity (β_art_ and pupillary dilation) in anticipation of the crash. These would hypothetically activate the network, including the anterior insula, and/or modulate functional connectivity (FC) between the anterior insula and other brain regions.

## Materials and Methods

### Experimental design

Study participants performed a simple reaction task while watching a driving video clip during an MRI scan. Figure 1 shows a schematic of the experimental trial.

**Figure 1.**
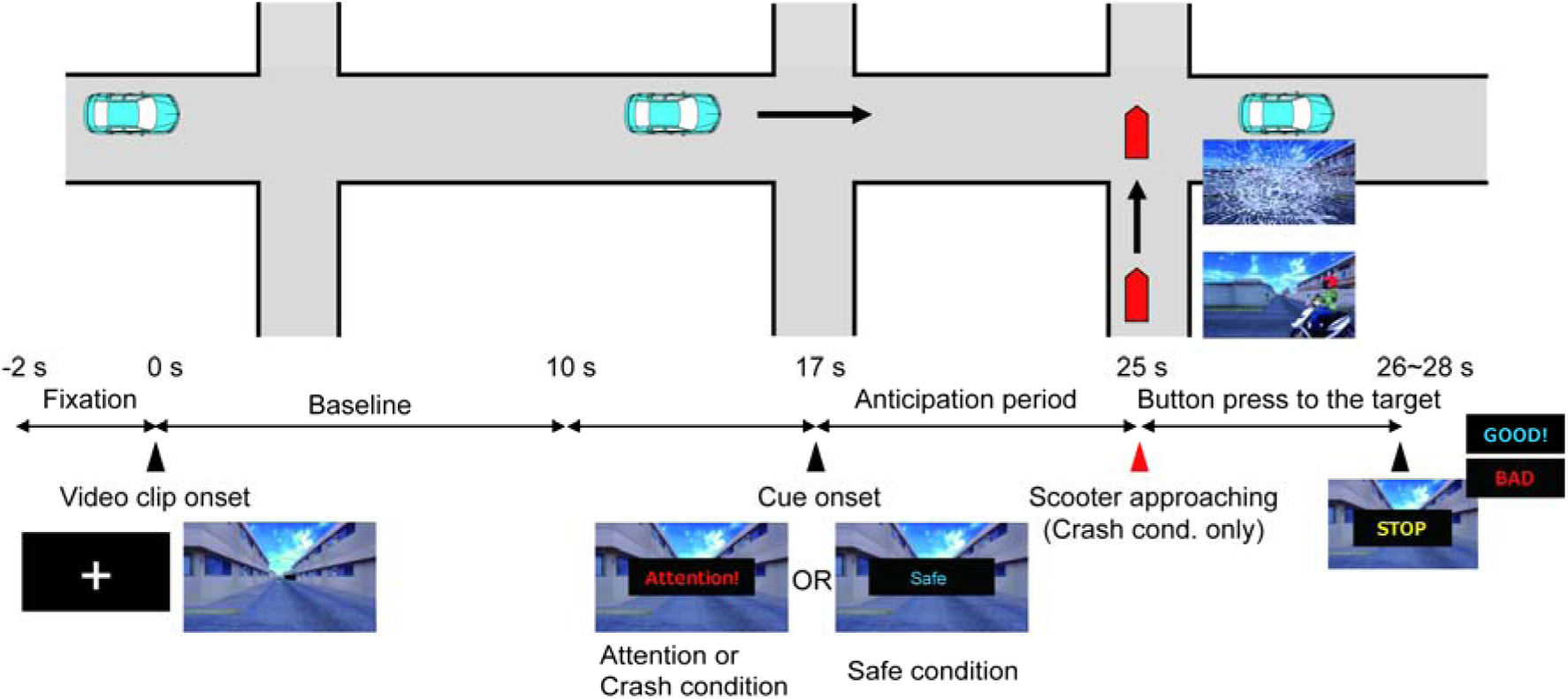
Schematic timeline of an experimental trial. In each trial, participants were presented with a first-person perspective driving video clip. At 17 s following the video clip onset, participants were presented with a word, either “Attention” or “Safe,” for 2 s as a cue to indicate whether a subsequent crash may occur (attention/crash condition) or not (safe condition). In the crash condition, a scooter approached and hit the vehicle at 25 s following the video clip onset. At the end of the video clip, the word “STOP” was presented for 1 s, and participants were asked to press a button as quickly as possible. They received feedback as to whether their response was performed within 1 s following the presentation of the word “STOP,” via presentation of a word, either “GOOD” or “BAD.”. We defined the 10-s period from the video clip onset as “baseline,” and the 8-s period from the cue onset as the “anticipation period.” See “Materials and Method” for details.

In the experiment, participants viewed a first-person perspective driving video clip generated by a driving simulator (D3sim, Mitsubishi Precision Co., Ltd., Japan). The video clip displayed a vehicle advancing through a narrow street for 26–28 s. At 17 s following the onset of the video clip, either the word “Attention” or “Safe” was superimposed for 2 s on the video clip. Among the 16 trials in which the word “Attention” appeared, six revealed a scooter approaching and hitting the vehicle, with the presentation of a broken windshield and a collision sound (crash condition). In contrast, no scooter approached the vehicle in the remaining 10 trials (attention condition) and in the 16 trials in which the word “Safe” appeared (safe condition). Before the experiment, the participants were informed of the potential scenarios after the presentation of each type of cue. At the end of each trial, the word “STOP” was displayed for 1 s, upon which the participants had to press a button with their right index finger, similar to a hard-braking event. The timing of the presentation of the word “STOP” was randomized within 26–28 s following the video clip onset. Participants were presented with the word “GOOD” or “BAD” as feedback to indicate whether they responded within 1 s from the presentation of the word “STOP.” We defined the 10-s period from the video clip onset as the “baseline” period and the 8-s period from the cue onset as the “anticipation” period.

After each trial, participants rated four items (anxiety, pleasantness, unpleasantness, and arousal), assessing their affective states during the anticipation period using a visual analog scale (VAS; Aitken 1969). For each item, the participants moved a cursor to indicate their rating (from 0 to 100) by pressing buttons with their middle and ring fingers. The experiment consisted of four sessions of eight trials each. Prior to the main trials, the participants performed three practice trials (one trial for each condition). Trait and state anxiety scores were obtained using the State-Trait Anxiety Inventory (STAI). Trait anxiety scores were obtained prior to the experiment. State anxiety scores were obtained before and after the experiment.

Visual stimuli were presented on an MRI-compatible, 32-inch liquid crystal display (NordicNeuroLab, Bergen, Norway) with a resolution of 1,920 × 1,080 pixels that subtended 30.4° × 17.4° visual angles. The participants perceived the visual stimuli through a mirror attached to the head coil. Auditory stimuli were presented with MRI-compatible, noise-canceling headphones (OptoActive, Optoacoustics Ltd., Or Yehuda, Israel). The volume of the auditory stimuli was adjusted to ensure that it was not too loud based on the participants’ self-reports during the practice trials.

### Participants

Thirty-four healthy adult participants completed this experiment (four women, all right-handed, aged 23.0 ± 2.12 [mean ± SD]). All participants had a driver’s license and drove a car at least once a week. According to self-report data, none of the participants had a history of mental disorders. All participants provided written informed consent prior to participating in the study. This study was approved by the Research Ethics Committee of Hiroshima University (approval number E-965-3).

### Analysis of behavioral data

The subjective rating data for each item were z-score normalized for each participant. To examine the differences in ratings among conditions and the occurrence of habituation, we performed a two-way repeated-measures analysis of variance (ANOVA) with factors of task condition and session for each item.

The reaction time (RT) data for each participant were averaged for each condition in the four sessions. To examine whether there were any differences in the mean RT, we performed a one-way repeated measures ANOVA with a factor of task condition. If any main effects or interactions were significant, we performed a post-hoc test using the modified Shaffer method.

### Acquisition and analyses of autonomic responses

To examine autonomic responses related to the participants’ anxiety, we measured the pupil area and blood pressure, and we performed ECG and PPG.

### Pupil area

The pupil area of the participants’ right eye was monitored using an MRI-compatible eye tracker (EyeLink 1000 Plus, SR Research, Osgoode, ON, Canada) at a sampling rate of 500 Hz. The pupil data were analyzed within the period from −2 s to 25 s of the onset (0 s) of the video clip. Data for one participant and those of six out of the remaining 132 sessions (33 participants × 4 sessions) were excluded from the statistical analysis owing to excessive eye blinks and measurement artifacts based on visual inspection. To avoid misrepresentation of a dataset, the data 200 ms before and after the eye blinks were replaced with “Not a Number” (NaN) since the data within this period were possibly distorted by eye blinks (Choe et al. 2016). To examine when a difference was statistically significant in the trial, we performed a paired *t*-test on the mean pupil areas between the conditions at each time point throughout the trial. Since we expected a difference in the autonomic responses between the attention and safe conditions during the anticipation period, and since the presentation of a cue itself could affect the autonomic responses right after the cue onset, we also calculated the mean pupil areas over the 5-s period from 3 s to 8 s following cue onset (20–25 s following video clip onset), and compared values between conditions using a paired *t*-test. We focused on the period before the crash (25 s following video clip onset); therefore, the crash and attention conditions were collapsed (crash/attention condition) for this analysis.

### Peripheral arterial stiffness

ECG signals were acquired from a three-lead ECG placed on the participant’s chest. A PPG was attached to the participant’s left index finger. Systolic and diastolic blood pressure values during a single heartbeat were measured using the CareTaker system (Biopac Systems, Goleta, CA, USA). A blood pressure cuff was attached to the participant’s left thumb. These physiological signals were recorded using a Biopac MP 150 system at a sampling rate of 1,000 Hz. Since data could not be obtained from one participant owing to a problem in the measurement device, the data of 33 participants were used for the statistical analysis. β_art_ was calculated for each heartbeat based on the following equation (Tsuji et al. 2021):

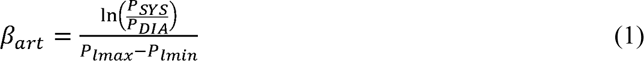

where *P_lmax_* and *P_lmin_* are the maximum and minimum values of the PPG during a heartbeat, respectively. *P_SYS_* and *P_DIA_* are systolic and diastolic blood pressures during a heartbeat. The ECG was used to determine the R–R interval for extracting the *P_SYS_*, *P_DIA_*, *P_lmax_*, and *P_lmin_* for each heartbeat. However, we could not robustly detect the R peaks in the ECG data obtained from most participants owing to MRI artifacts. Therefore, we used the inverted PPG waveforms for each participant and segmented the waveform between two adjacent peaks to determine each heartbeat.

To compare the β_art_ between the conditions, we calculated the ratio of the mean β_art_ averaged over the 5-s period from 3 to 8 s following cue onset (20 to 25 s following video clip onset) to the mean β_art_ averaged over the baseline, and subsequently performed a paired *t*-test.

### MRI data acquisition

MRI data were acquired using a 3.0T MRI scanner (Siemens Magnetom Verio, Siemens AG, Munich, Germany). The functional images were acquired using an echo-planar, T2*-weighted, multiband gradient echo sequence with the following parameters: repetition time (TR), 1,000 ms; echo time (TE), 30.00 ms; voxel size, 3.0 × 3.0 × 3.2 mm; 42 slices; slice thickness, 3.2 mm; field of view (FOV), 192 mm; flip angle, 80°; and acceleration factor, 3. The structural image of each participant was acquired using T1-weighted, three-dimensional magnetization-prepared rapid gradient-echo imaging (MPRAGE) with the following parameters: TR, 2,300 ms; TE, 2.98 ms; voxel size, 1.0 × 1.0 × 1.0 mm; flip angle, 9°; and FOV, 256 mm.

### MRI data analysis

MRI data preprocessing and statistical analyses were performed using SPM12 software (Wellcome Department of Cognitive Neurology, London, UK; www.fil.ion.ucl.ac.uk/spm). The first 10 echo-planar images (EPIs) were discarded to permit T1 equilibration effects. In order to account for the correction of head movement, the remaining volumes were spatially realigned to the first of the volumes and realigned to the mean of all the images. T1-weighted structural images were co-registered with the first EPI of the remaining EPIs after discarding the first 10 EPIs. The co-registered structural images were spatially normalized to the Montreal Neurological Institute template. The parameters derived from this normalization process were subsequently applied to each EPI. The normalized EPIs were spatially smoothed using an 8-mm full width at half maximum Gaussian kernel.

Voxel-based statistical analysis of the pre-processed EPIs was performed using a general linear model (GLM). The blood oxygenation level-dependent (BOLD) response was related to the baseline and anticipation periods, and the crash events were modeled as a boxcar function for the onset and duration of each event (10 s for the baseline, 8 s for the anticipation period, and 0 s for the crash event). Each resulting time series for each event was convolved with a canonical hemodynamic response function, which was subsequently used as a regressor. Six head motion parameters, derived from the realignment process, were also used as regressors to reduce motion-related artifacts. To eliminate low-frequency drifts, a high-pass filter with a 128-s cut-off period was applied to the fMRI time series. Serial autocorrelations between scans were corrected using a first-order autoregressive model.

Regression coefficients for each event were computed for each participant using a fixed-effects model. We created contrast images for the anticipation periods relative to the baseline in each of the crash, attention, and safe conditions (anticipation_crash > baseline; anticipation_attention > baseline; anticipation_safe > baseline) for the brain activity for each event. In addition, we created contrast images to compare the brain activity during the anticipation period between the attention and safe conditions (anticipation_attention > anticipation_safe; anticiaption_safe > anticipation_attention). We also created contrast images for the crash events relative to the baseline. These contrast images were subjected to group analysis using a random-effects model with a one-sample *t*-test. In the group analysis, we set the statistical threshold at an uncorrected *p* < 0.001 at the voxel level and at a family-wise error (FWE)-corrected *p* < 0.05 at the cluster level. We excluded the crash condition (i.e., the condition in which the cue, “Attention,” was followed by a car crash) from the main analyses (for the common brain activity during the anticipation period, the difference in brain activity during the anticipation period between the conditions, and the parametric modulation analyses), based on the following reasons: (1) the crash condition contained the presentation of a broken windshield and a collision sound, unlike the attention and safe conditions; (2) since subjective ratings were obtained after each trial, it is possible that the additional aversive auditory-visual information affected the subjective ratings in the crash condition.

We conducted parametric modulation analyses to examine subjective anxiety-related brain activity. For parametric modulation of subjective anxiety, we used regressors for the anticipation period in the trials under both attention and safe conditions, and z-score normalized scores of anxiety in each corresponding trial as the parametric modulator.

For autonomic response-related brain activity, we conducted parametric modulation analyses using β_art_ and the pupil area. In these analyses, trial-by-trial values of the ratio of mean β_art_ and pupil area were used. These were calculated by averaging data over the 5-s period (from 3 to 8 s) following the cue onset to those averaged over the baseline. We created parametric modulation regressors using β_art_ and the pupil area for the anticipation period in the attention and safe conditions.

### Autonomic response-related functional connectivity analysis

To examine the pupil- and β_art_-related brain network, we examined FC between the anterior insula and its adjacent regions (as seed regions) and the rest of the brain regions by conducting a generalized form of context-dependent psychophysiological interactions (gPPI) using the CONN toolbox (Whitfield-Gabrieli and Nieto-Castanon 2012). For gPPI analyses, we created GLMs, including a regressor for the anticipation period in trials under both attention and safe conditions. The processed GLM for each participant was imported to the gPPI model in the CONN toolbox. For each participant, we created a separate gPPI model for pupil area and β_art_ parametric modulators. As first-level covariates, we created gPPI regressors using parametric-modulation time-series based on trial-by-trial values of the ratio of the mean β_art_ and pupil area averaged over the 5-s period from 3 to 8 s following the cue onset to those averaged over the baseline. Based on our hypothesis, we defined seed regions as the anterior insula (dorsal and ventral agranular insula) and the anatomical regions in the cluster, including the anterior insula activated in the contrast condition (attention > safe). We used the human Brainnetome Atlas (Fan et al. 2016) to define these anatomical regions of interest. In the second-level analysis, seed-based connectivity maps of Fisher-transformed correlation coefficients between BOLD time-series of each seed region and each individual voxel were entered into a one-sample *t*-test to examine the brain regions showing significant pupil-and β_art_-related FC with each seed region. We applied a statistical threshold at an uncorrected *p* < 0.001 at the voxel level and a false discovery rate (FDR)-corrected *p* < 0.05 at the cluster level.

## Results

### Behavioral data

#### Subjective ratings

Figure 2 shows the results of subjective ratings. Regarding subjective rating of anxiety, a repeated-measures two-way ANOVA with factors of task conditions (crash, attention, and safe) and session (sessions 1, 2, 3, and 4) revealed a significant main effect of task conditions (*F* [1.84, 60.83] = 316.596, partial η^2^ = 0.906, *p*_corrected_ < 0.001; Chi-Muller correction for non-sphericity was applied). Post-hoc analyses using the modified Shaffer method revealed that the participants rated subjective anxiety highest in the crash condition, followed by the attention condition and the safe condition (all *t*s [33] > 4.281, all *p*s < 0.001). In contrast, we observed no main effect of session (*F* [2.40, 79.08] = 0.423, partial η^2^ = 0.013, *p*_corrected_ = 0.693) or interaction (*F* [5.35, 176.41] = 0.786, partial η^2^ = 0.023, *p*_corrected_ = 0.569). Considering the unpleasantness and arousal, repeated-measures two-way ANOVA also revealed a significant main effect of condition (unpleasantness: *F* [2, 66] = 304.478, partial η^2^ = 0.902, *p*_corrected_ < 0.001; arousal: *F* [2, 66] = 156.673, partial η^2^ = 0.826, *p*_corrected_ < 0.001); however, no main effect of session (unpleasantness: *F* [2.64, 86.96] = 1.004, partial η^2^ = 0.03, *p*_corrected_ = 0.388; arousal: *F* [2.54, 83.78] = 0.696, partial η^2^ = 0.021, *p*_corrected_ = 0.534) or interaction (unpleasantness: *F* [4.93, 162.75] = 0.898, partial η^2^ = 0.026, *p*_corrected_ = 0.483; arousal: *F* [4.99, 164.57] = 1.316, partial η^2^ = 0.038, *p*_corrected_ = 0.260) was observed. Post-hoc analyses again replicated the result for anxiety (unpleasantness: all *t*s [33] > 10.207, *p* < 0.001; arousal: all *t*s [33] > 3.002, all *p*s < 0.01).

**Fig. 2.**
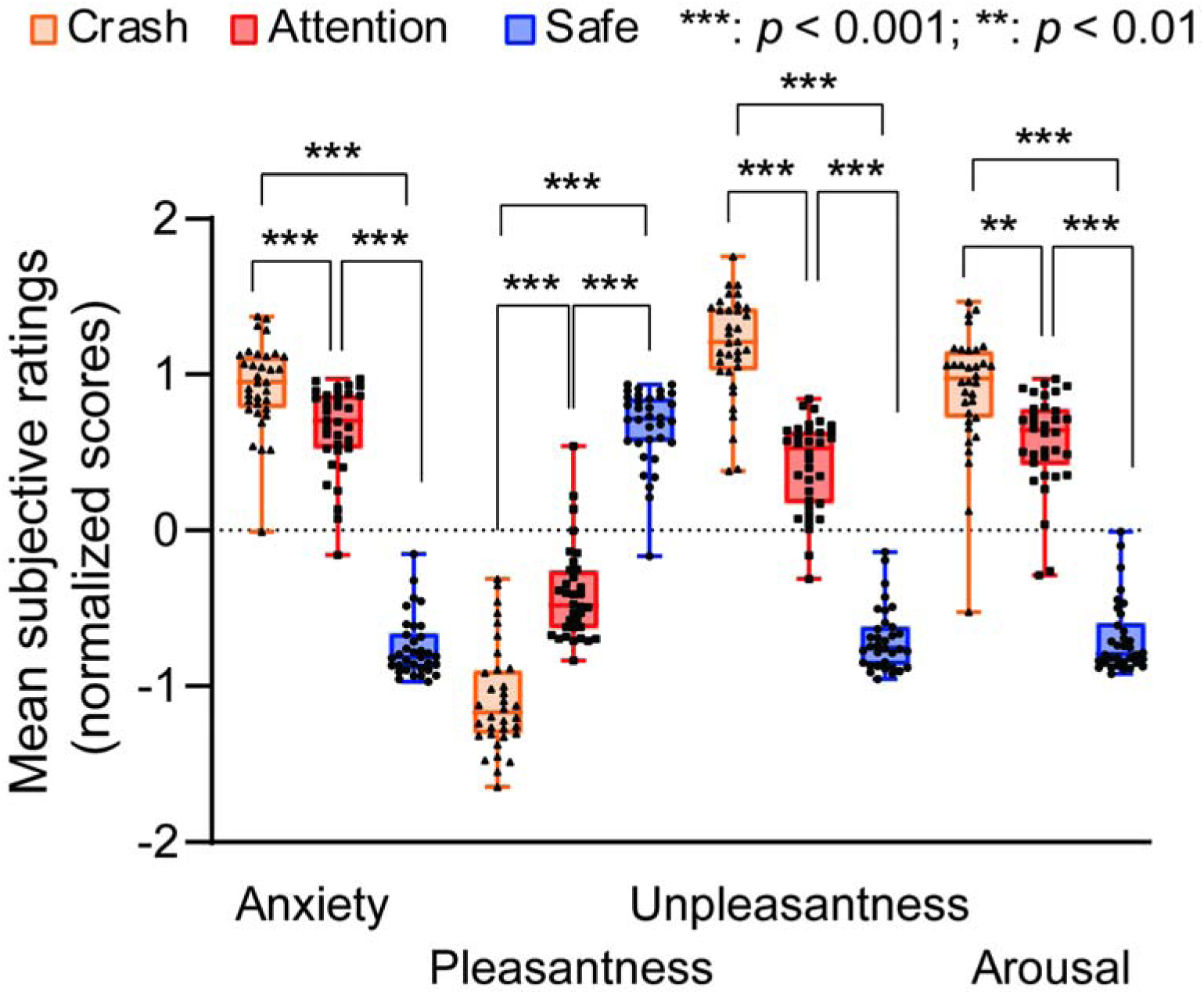
The results of subjective ratings (N = 34).

Considering pleasantness, a repeated-measures two-way ANOVA again revealed a significant main effect of condition (*F* [1.94, 63.91] = 213.201, partial η^2^ = 0.866, *p*_corrected_ < 0.001). Post-hoc analyses revealed that pleasantness was rated highest in the safe condition, followed by the attention condition and the crash condition (all *t*s [33] > 9.249, all *p*s < 0.001). We observed no main effect of session (*F* [3, 99] = 1.005, partial η^2^ = 0.03, *p*_corrected_ = 0.394) and no significant interaction (*F* [4.89, 161.29] = 2.098, partial η^2^ = 0.06, *p*_corrected_ = 0.07).

The mean scores of STAI-trait, and STAI-state before (STAI-state pre) and after the experiment (STAI-state post) were as follows: STAI-trait, 42.5 ± 7.50; STAI-state pre, 35.29 ± 7.50; and STAI-state post, 34.82 ± 6.80. We conducted correlation analyses between all pairs of the mean subjective ratings for each participant for each item and the STAI score (STAI-trait, STAI-state pre, and STAI-state post). No significant correlation was observed between any of the pairs (all *p*s > 0.10).

#### Reaction times

Figure 3 shows the mean RT for each condition. A one-way ANOVA on the mean RTs with a factor of task condition revealed a significant main effect (*F* [1.73, 57.07] = 17.793, partial η^2^ = 0.350, *p*_corrected_ < 0.001). Post-hoc analyses using the modified Shaffer method revealed that the mean RT for the crash condition was faster than that for the attention and safe conditions (*t*s [33] < −4.060, *p*s < 0.001). However, we found no significant difference between the attention and safe conditions (*t* [33] = −1.070, *p* = 0.292).

**Fig. 3.**
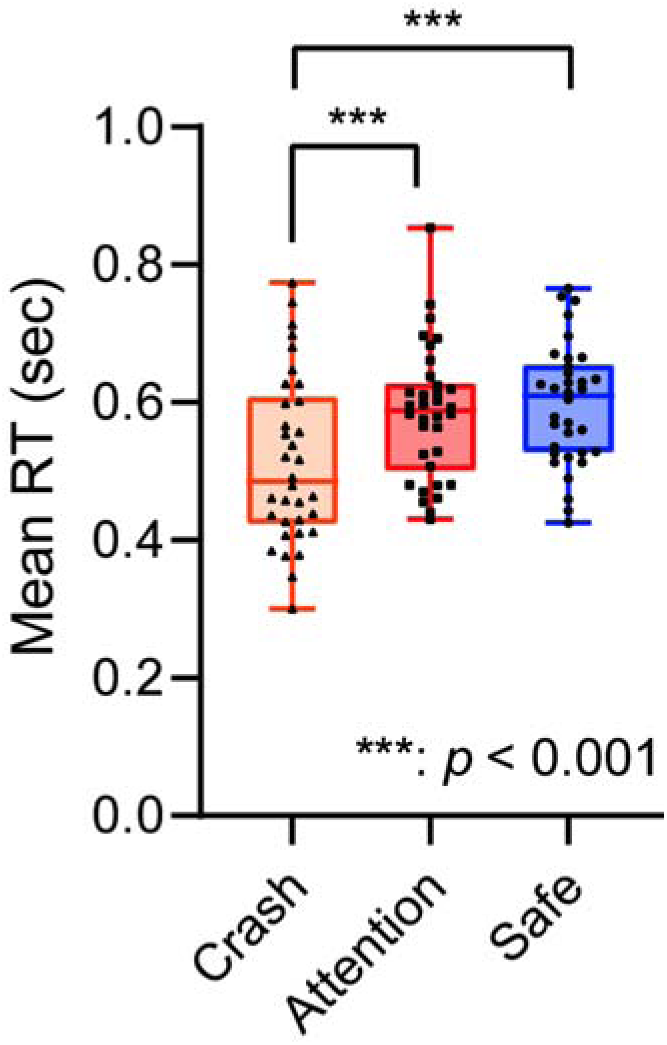
Mean reaction time (N = 34).

### Autonomic responses

#### Pupil dilation

Figure 4 shows the mean time series of all the participants’ pupil responses during the task for each condition. For data at each time point, we performed an FDR-corrected paired *t*-test. This revealed significantly more pupil dilation in the crash/attention conditions than in the safe condition at a threshold of *p*_FDR_ < 0.05, mainly in the period from approximately 1 s following the cue onset (i.e., 18 s following the video clip onset) to the end of the trial. The gray areas in Figure 4 represent the time points showing significant differences between the conditions. This indicates that significant differences were robustly observed during the period from 3 to 8 s following cue onset. To examine the statistical difference between conditions during this period, we performed a paired *t*-test between the mean pupil areas in the crash/attention and safe conditions in the period from 3 to 8 s following the cue onset. This again revealed significantly more pupil dilation in the crash/attention condition than in the safe condition (*t* [33] = 4.4023, *p* < 0.001).

**Fig. 4.**
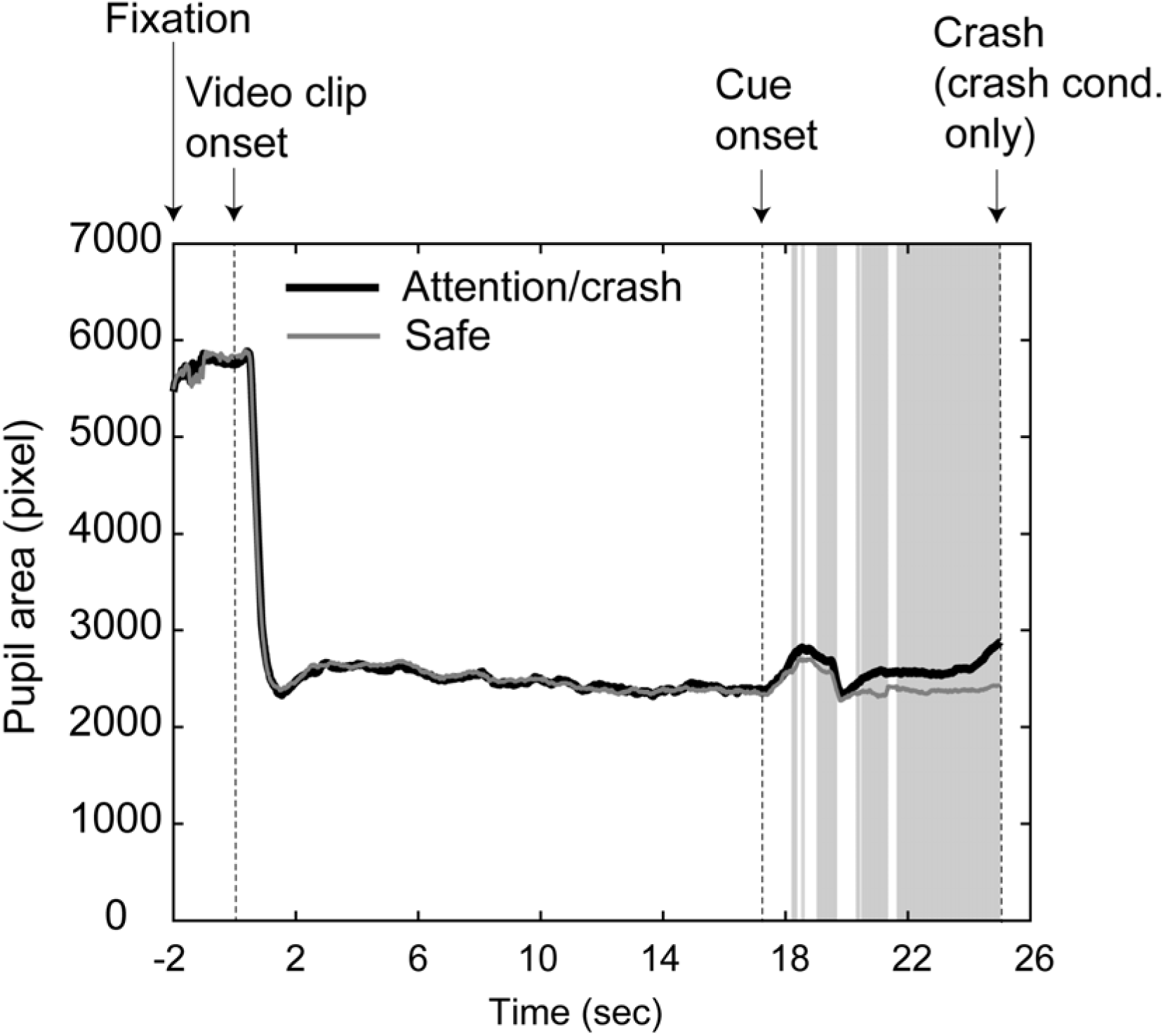
The mean time-series of pupil areas for attention/crash and safe conditions. The gray areas represent the time points showing significant differences between conditions (see text).

To examine the relationship between the pupil dilation and the subjective ratings of anxiety, we conducted a correlation analysis using the difference between the mean rating scores of anxiety in the attention and safe conditions for each participant, and the difference between the mean pupil areas in the crash/attention and safe conditions for the period from 3 to 8 s following the cue onset. The analysis revealed that the pupil dilation and the anxiety ratings were not significantly correlated (*r* = 0.313, *t* [31] = 1.834, *p* = 0.076, Supplementary Figure S1). For the other items, we also observed no significant correlation (arousal: *r* = 0.318, *t* [31] = 1.865, *p* = 0.072; pleasantness: *r* = 0.270, *t* [31] = 1.564, *p* = 0.128; unpleasantness: *r* = 0.149, *t* [31] = 0.836, *p* = 0.409).

#### Peripheral arterial stiffness

Figure 5 shows the mean time series of all the participants’ β_art_ during the task for each condition. To examine the significance of the increase in β_art_ following cue onset, we compared the ratio of the mean value of β_art_ during the 5-s period from 3 to 8 s following the cue onset to the mean value over the baseline between the attention and safe conditions using a paired *t*-test. This revealed non-significant differences in β_art_ between the attention condition and the safe condition (*t* [32] = 1.80, *p* = 0.08).

**Fig. 5.**
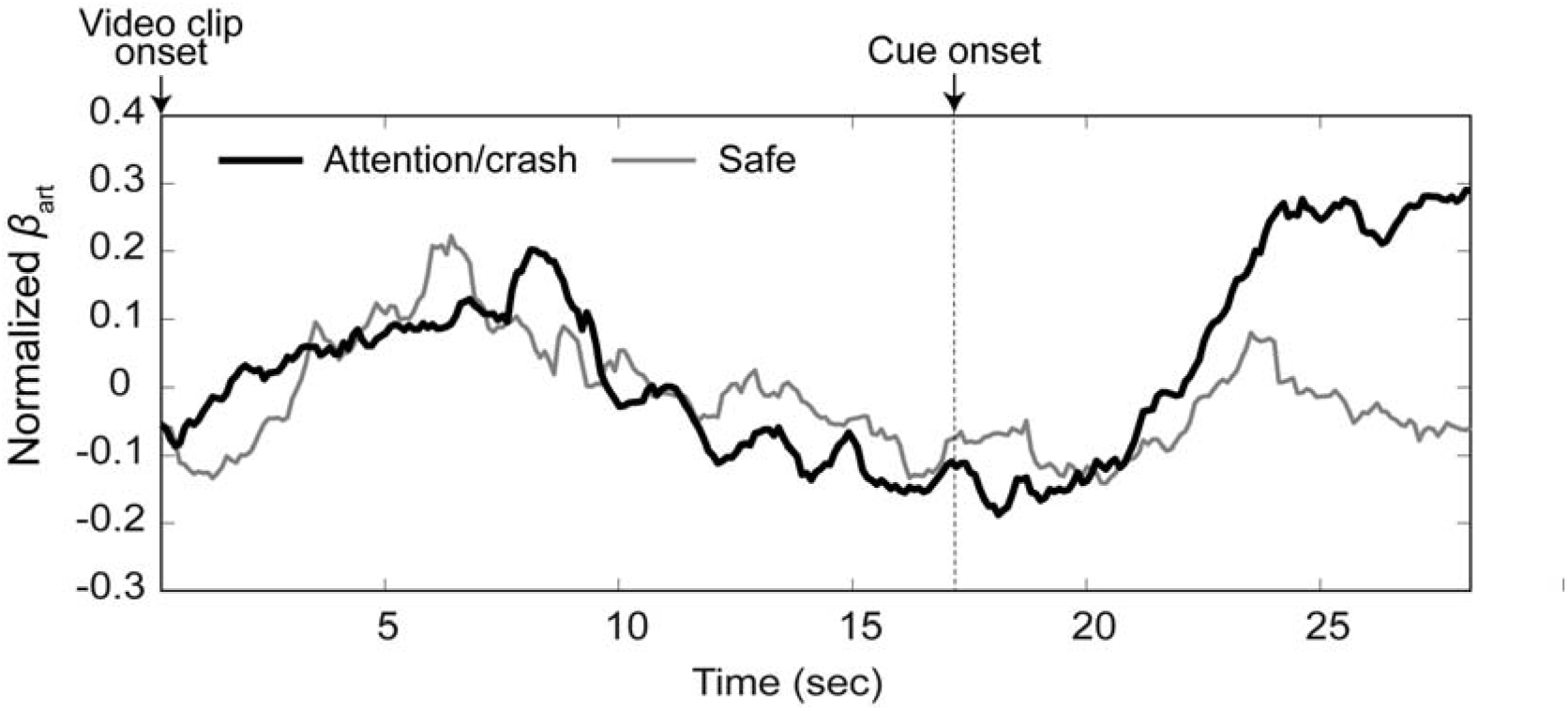
The mean time-series of peripheral arterial stiffness (β_art_) for attention/crash and safe conditions.

### fMRI data

#### Brain activity during the anticipation period

To examine the common brain activity during the anticipation period relative to the baseline in both the attention and safe conditions, we conducted a conjunction analysis. This revealed significant activations in the bilateral front-parietal cortices, dorsal anterior cingulate cortex, and bilateral anterior insulae (Figure 6, Table 1).

**Fig. 6.**
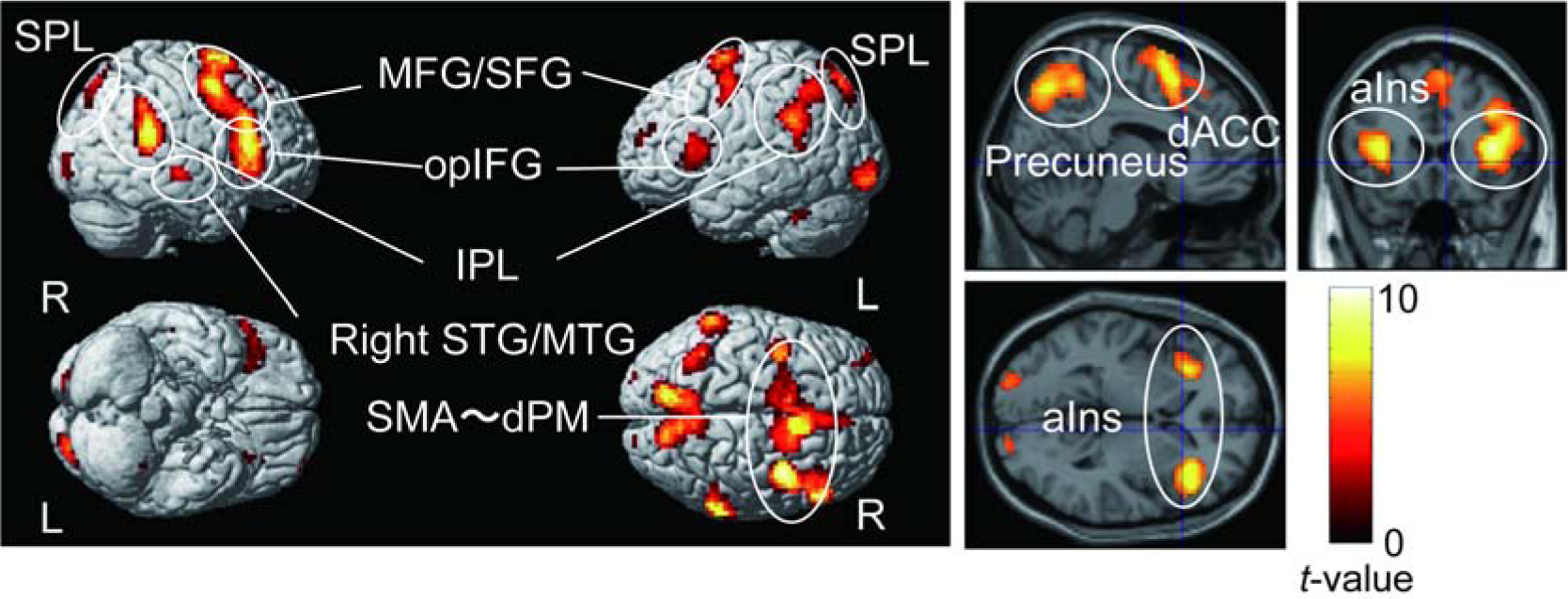
Brain regions commonly activated in the attention and safe conditions revealed by a conjunction analysis. SPL, superior parietal lobule; MFG, middle frontal gyrus; SFG, superior frontal gyrus; opIFG, inferior frontal gyrus, opercular part; IPL, inferior parietal lobule; SMA, supplementary motor area; dPM, dorsal premotor cortex; dACC, dorsal anterior cingulate cortex; aIns, anterior insula

**Table 1.** Common brain regions activated in both the attention and safe conditions during the anticipation period based on the result of a conjunction analysis.

| Anatomical region | Cluster size | MNI coordinates (mm) |  |  | T-value |
| --- | --- | --- | --- | --- | --- |
|  |  | x | y | z |  |
| Left lingual gyrus | 4,299 | -12 | -88 | -9 | 18.67 |
| Right calcarine gyrus |  | 12 | -88 | -3 | 17.87 |
| Right lingual gyrus |  | 21 | -76 | -12 | 15.84 |
| Right superior frontal gyrus | 1201 | 24 | -1 | 61 | 9.34 |
| Left superior frontal gyrus |  | -24 | -4 | 68 | 8.41 |
| Left supplementary motor area |  | -6 | 11 | 52 | 7.39 |
| Right inferior frontal gyrus,<br>triangular part | 116 | 60 | 23 | 16 | 5.35 |
| Right inferior frontal gyrus,<br>triangular part/anterior insula |  | 39 | 23 | 13 | 4.45 |
| Left inferior frontal gyrus,<br>opercular part/left anterior insula | 100 | -45 | 14 | 10 | 5.15 |
| Left inferior frontal gyrus,<br>opercular part/left anterior insula |  | -36 | 17 | 10 | 4.70 |

Uncorrected $p < 0.001$ at peak level, family-wise error-corrected $p < 0.05$ at the cluster level. MNI,
Montreal Neurological Institute

To examine the brain regions related to each condition, we compared brain activation between the attention and safe conditions during the anticipation period. In the attention condition, the right anterior insula and the subcortical regions, including the PAG, thalamus, and BNST were more active than in the safe condition (Figure 7, Table 2). We confirmed that the activated cluster had overlaps with the atlas of the PAG (Keuken et al. 2017) and BNST (Torrisi et al. 2015, Figure 7B). In contrast, the areas from the parietal operculum, including the secondary somatosensory area (SII) to the posterior insula, and the precuneus were less active in the attention condition than in the safe condition. Although these areas were deactivated under all conditions, deactivation was more evident in the attention condition than in the safe condition (Figure 8, Table 3).

**Fig. 7.**
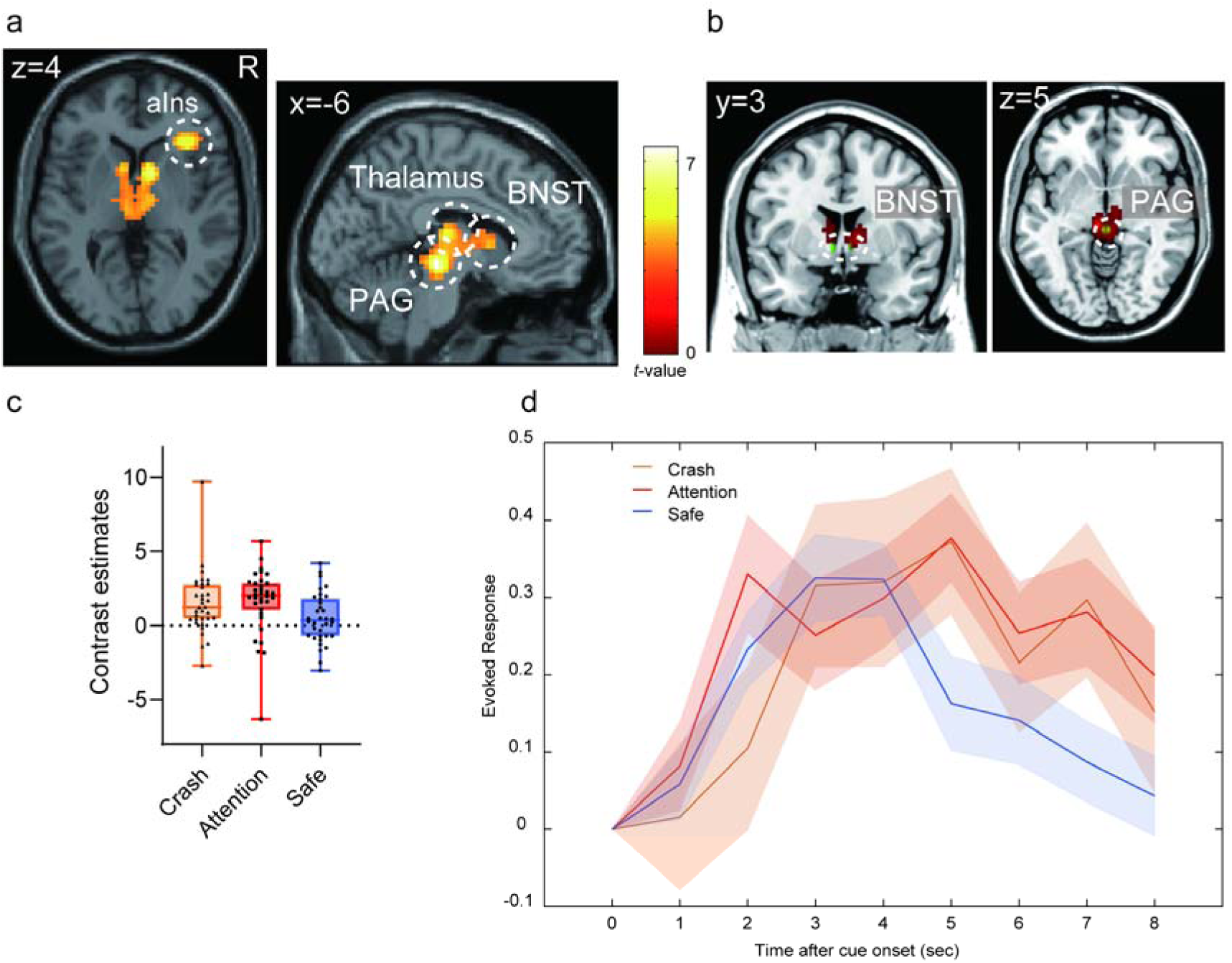
(A) Brain regions showing more activation in the attention condition than in the safe condition. (B) The overlap between the activated cluster (red) and the anatomical atlas of the BNST (Torrisi et al. 2015) and PAG (Keuken et al. 2017) (green). (C) Contrast estimates of the peak voxel in the activated cluster in the right anterior insula for each condition. (D) The time course for BOLD signal change of the peak voxel in the right anterior insula for each condition during the anticipation period. Each line represents the mean event-related BOLD response over all participants for each condition with the standard error of the mean as represented by the area with the corresponding color. The plot was created by using the rfxplot Toolbox (Gläscher 2009).

**Fig. 8.**
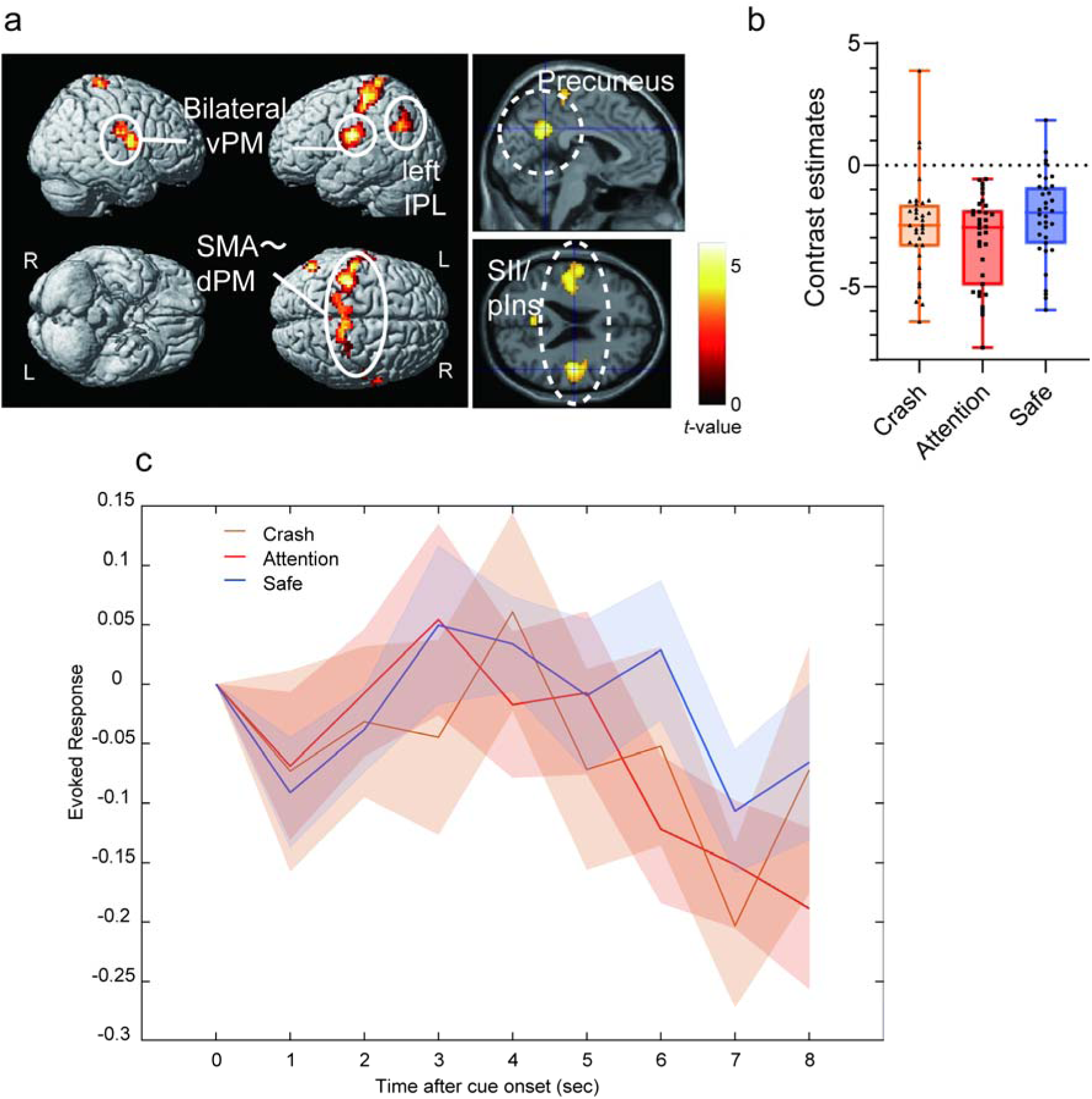
(A) Brain regions showing more activation in the safe condition than in the attention condition. (B) Contrast estimates of the peak voxel in the activated cluster in the right SII/posterior insula for each condition. (C) The time course for blood oxygenation level-dependent (BOLD) signal change of the peak voxel in the right SII/posterior insula for each condition during the anticipation period. Each line represents the mean event-related BOLD response over all participants for each condition with the standard error of the mean as represented by the area with the corresponding color. The plot was created by using the rfxplot Toolbox (Gläscher 2009).

**Table 2.** Brain regions significantly more activated in the attention condition than in the safe condition

| Anatomical region | Cluster<br>size | MNI coordinates<br>(mm) |  |  | *T*-value |
| --- | --- | --- | --- | --- | --- |
|  |  | x | y | Z |  |
| Left red nucleus | 516 | -6 | -25 | -9 | 7.53 |
| Right caudate head |  | 12 | 5 | 4 | 6.31 |
| Left medial dorsal thalamus |  | -9 | -19 | 13 | 5.97 |
| Right inferior frontal gyrus,<br>triangular part/anterior insula | 105 | 36 | 26 | 7 | 5.96 |
| Right inferior frontal gyrus,<br>triangular part |  | 48 | 29 | 16 | 3.76 |

5 Uncorrected $p < 0.001$ at peak level, family-wise error-corrected $p < 0.05$ at the cluster level. MNI,
6 Montreal Neurological Institute

**Table 3.** Brain regions significantly more activated in the safe condition than in the attention condition

| Anatomical region | Cluster<br>size | MNI coordinates<br>(mm) |  |  | *T*-value |
| --- | --- | --- | --- | --- | --- |
|  |  | x | y | z |  |
| Right Rolandic operculum | 303 | 48 | -13 | 23 | 5.70 |
| Right Rolandic operculum |  | 60 | 2 | 13 | 5.26 |
| Right posterior insula |  | 39 | -10 | 16 | 4.99 |
| Left posterior cingulate cortex | 110 | -3 | -52 | 32 | 5.10 |
| Left postcentral gyrus | 307 | -48 | -19 | 26 | 5.06 |
| Left posterior insula |  | -39 | -13 | 13 | 4.95 |
| Left postcentral gyrus |  | -57 | -7 | 20 | 4.54 |
| Left inferior parietal cortex | 500 | -54 | -22 | 48 | 4.92 |
| Left postcentral gyrus |  | -42 | -34 | 61 | 4.74 |
| Right paracentral lobule |  | 6 | -31 | 64 | 4.74 |
| Left angular gyrus | 99 | -57 | -67 | 29 | 4.85 |
| Left angular gyrus |  | -45 | -55 | 29 | 4.28 |
| Left angular gyrus | 303 | -51 | -67 | 42 | 3.90 |

Uncorrected $p < 0.001$ at peak level, family-wise error-corrected $p < 0.05$ at the cluster level. MNI,
Montreal Neurological Institute

Parametric modulation using anxiety rating scores, including attention and safe conditions, revealed that activations in the thalamus and PAG were positively correlated with the anxiety ratings. In contrast, we found no brain regions showing a significant negative correlation with the anxiety ratings at a voxel-level threshold of uncorrected *p* < 0.001 or a cluster-level FWE-corrected *p* < 0.05; however, at a liberal threshold, the SII and posterior insula showed a negative correlation at a voxel-level uncorrected *p* < 0.005 (cluster-corrected FWE-corrected *p* = 0.066; Figure 9, Table 4).

**Fig. 9.**
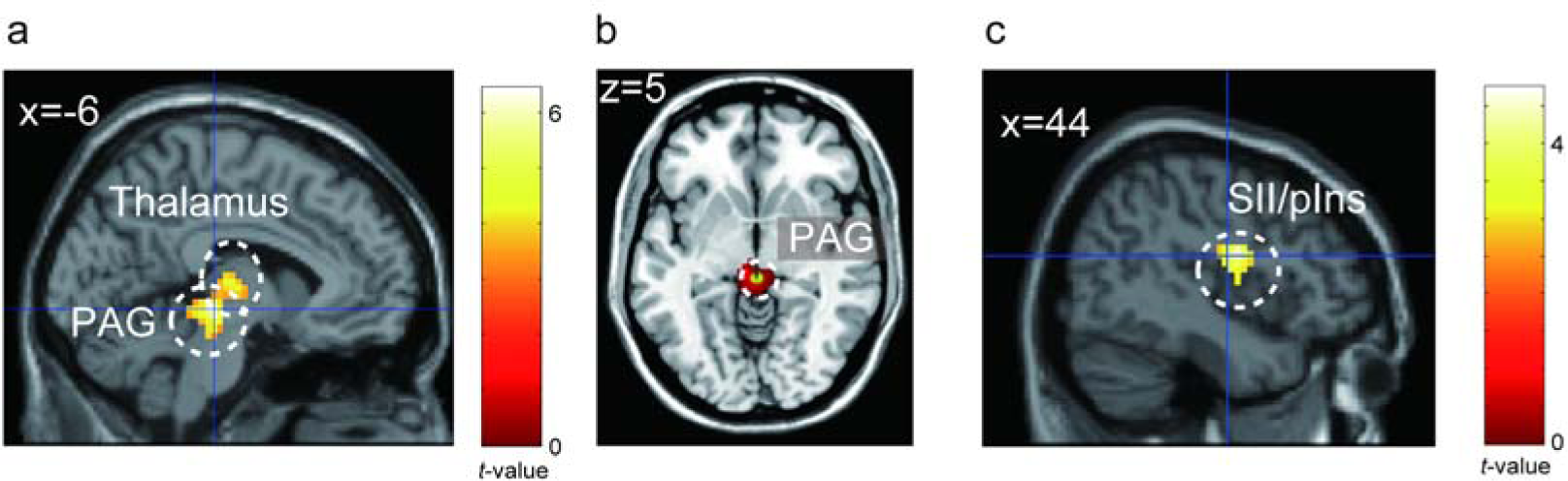
Brain activation correlated with the anxiety rating scores. (A) Brain regions positively correlated with subjective anxiety (B) The overlap between the activated cluster (red) and the anatomical atlas of the PAG (Keuken et al. 2017) (green). (C) Brain regions negatively correlated with subjective anxiety at a liberal threshold (voxel-level uncorrected *p* < 0.005 and cluster-level FWE-corrected *p* = 0.066). PAG, periaqueductal gray; BNST, bed nucleus of the stria terminalis; SII, secondary somatosensory area; pIns, posterior insula.

**Table 4.** (A) Brain regions showing activation positively correlated with subjective ratings of anxiety

| Anatomical region | Cluster | MNI coordinates |  |  | *T*-value |
| --- | --- | --- | --- | --- | --- |
|  | size | (mm) |  |  |  |
|  |  | x | y | z |  |
| Left red nucleus | 199 | -6 | -25 | -6 | 6.43 |
| Left medial dorsal thalamus |  | -3 | -19 | 4 | 4.49 |
| Left medial dorsal thalamus |  | -9 | -19 | 10 | 4.46 |

| Anatomical region | Cluster | MNI coordinates |  |  | *T*-value |
| --- | --- | --- | --- | --- | --- |
|  | size | (mm) |  |  |  |
|  |  | x | y | z |  |
| Right Rolandic operculum | 194 | 48 | -4 | 20 | 4.74 |
| Right Rolandic operculum |  | 51 | -13 | 23 | 3.82 |
| Right posterior insula |  | 42 | -4 | 7 | 3.56 |

6 Uncorrected $p < 0.005$ at peak level. MNI, Montreal Neurological Institute

On the other hand, parametric modulation using physiological signals did not reveal any significant activation for either pupil areas or β_art_. aIns, anterior insula; PAG, periaqueductal gray; BNST, bed nucleus of the stria terminalis; BOLD, blood oxygenation level-dependent.

vPM, ventral premotor cortex; IPL, inferior parietal lobule; SMA, supplementary motor area; dPM, dorsal premotor cortex; SII, secondary somatosensory area; pIns, posterior insula.

#### Brain activity related to the crash

The contrast in the crash event versus baseline showed extensive brain activation; therefore, we reported the brain regions with a voxel-level FWE-corrected *p* < 0.05. The activated regions can be classified into approximately three large clusters (Figure 10A, Table 5): (1) a posterior midline cluster including the precuneus, cerebellum, and thalamus; (2) a frontal midline cluster including the supplementary motor area and ACC, and (3) a fronto-temporoparietal cluster including the anterior insula, inferior frontal cortex, superior temporal cortices, parietal operculum, and posterior insula.

**Fig. 10.**
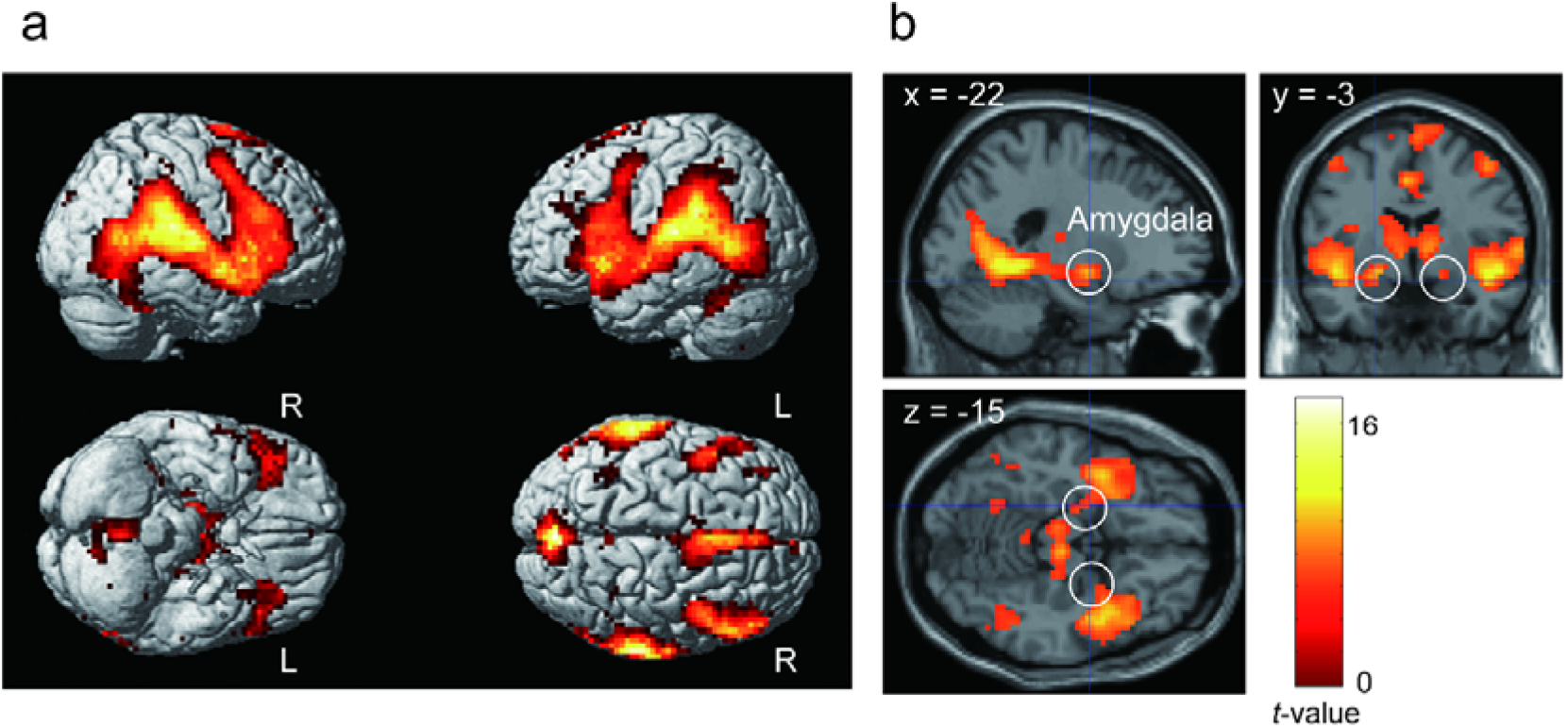
(A) Brain regions significantly activated in the contrast of crash > baseline. (B) The amygdala activation observed in the fronto-temporoparietal cluster.

**Table 5.** Brain regions significantly activated in the contrast crash > baseline

| Anatomical region | Cluster<br>size | MNI coordinates<br>(mm) |  |  | T-value |
| --- | --- | --- | --- | --- | --- |
|  |  | x | y | z |  |
| Left thalamus (medial pulvinar) | 11,381 | -9 | -25 | -3 | 17.48 |
| Right thalamus (medial geniculate nucleus) |  | 12 | -25 | -6 | 17.02 |
| Right lingual gyrus |  | 15 | -49 | -3 | 15.97 |
| Right pregenual ACC | 1,561 | 9 | 44 | 20 | 11.21 |
| Left superior medial frontal gyrus |  | 3 | 17 | 42 | 10.67 |
| Left supragenual ACC |  | -6 | 26 | 29 | 10.53 |
| Cerebellum vermis | 110 | 3 | -58 | -41 | 8.42 |
| Left inferior parietal lobule | 43 | -33 | -52 | 45 | 8.04 |
| Right paracentral lobule | 37 | 12 | -37 | 55 | 6.39 |
| Right precuneus |  | 12 | -46 | 55 | 6.20 |
| Right precuneus |  | 6 | -52 | 55 | 5.98 |
| Left cerebellum (VIII) | 35 | -12 | -70 | -48 | 6.27 |
| Cerebellum vermis |  | 6 | -73 | -41 | 6.16 |
| Left cerebellum crus |  | -9 | -76 | -35 | 5.92 |
| Left precuneus | 5 | -12 | -49 | 45 | 6.19 |
| Right inferior parietal lobule | 8 | 30 | -46 | 52 | 6.14 |
| Right inferior parietal lobule |  | 36 | -49 | 45 | 6.05 |
| Left middle frontal gyrus | 23 | -39 | 38 | 26 | 6.11 |
| Left middle frontal gyrus |  | -36 | 29 | 36 | 5.92 |
| Left middle frontal gyrus |  | -36 | 41 | 36 | 5.68 |
| Right superior parietal lobule | 1 | 54 | -43 | 58 | 5.74 |
| Left cerebellum (VIII) | 1 | -30 | -64 | -54 | 5.68 |
| Right inferior parietal lobule | 1 | 45 | -52 | 42 | 5.66 |

Family-wise error-corrected $p < 0.05$ at the peak level. MNI, Montreal Neurological Institute; ACC, anterior cingulate cortex

This fronto-temporoparietal cluster included the amygdala (Figure 10B).

#### Autonomic response-related functional connectivity

Based on the brain activity in the contrast condition (attention > safe), we conducted gPPI analyses using the right anterior insula (dorsal and ventral agranular insula) and the brain regions including the anterior insula cluster, such as the right middle frontal gyrus (MFG) ventral area (Brodmann area [BA] 9/46), right inferior frontal sulcus, right caudal inferior frontal gyrus (IFG) (BA45), right rostral IFG (BA45), and right opercular IFG (BA44) as seed regions.

Regarding the pupil area, although we observed no brain region showing significant FC for the right anterior insula as a seed region, we found brain regions showing significant pupil-related FCs with the brain areas adjacent to the anterior insula: the right MFG and the right rostral IFG (BA45). Moreover, the left cerebellum/brainstem showed a significant pupil-related FC with the right MFG, and the left visual cortex showed a significant pupil-related FC with the right rostral IFG (Figure 11A–B, Table 6).

**Fig. 11.**
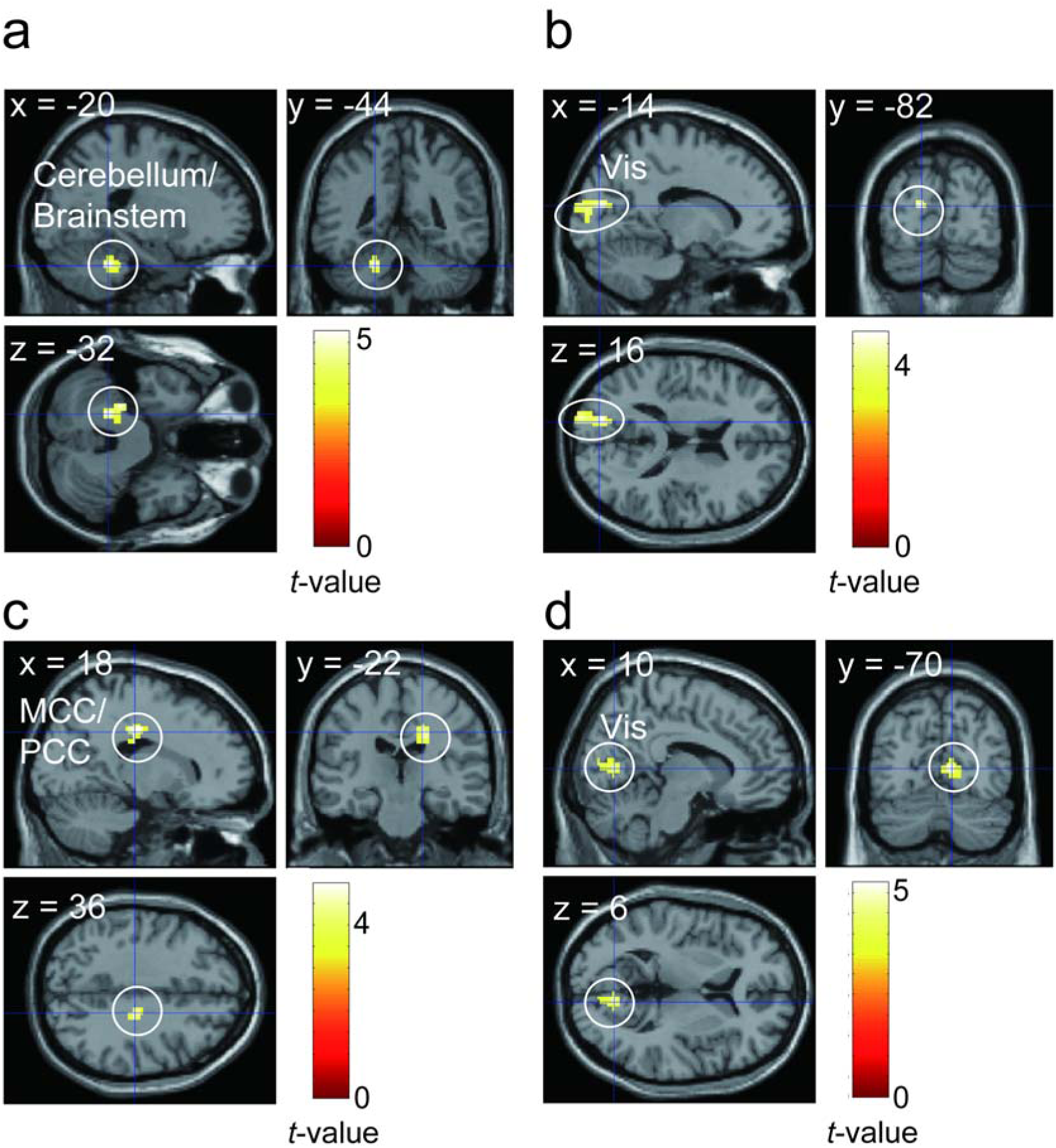
(A–B) Brain regions showing pupil-related functional connectivity. (A) Brain regions showing pupil-related functional connectivity with the right middle frontal gyrus as seed region. (B) Brain regions showing pupil-related functional connectivity with the right rostral inferior frontal gyrus as seed region. (C–D) Brain regions showing peripheral arterial stiffness (β_art_)-related functional connectivity. (C) Brain regions showing β_art_-related functional connectivity with the right dorsal agranular insula as seed region. (D) Brain regions showing β_art_-related functional connectivity with the right ventral agranular insula as seed region. Vis, visual cortex; MCC, middle cingulate cortex; PCC, posterior cingulate cortex.

**Table 6.** Brain regions showing pupil-related functional connectivity

(A) Brain regions showing pupil-related functional connectivity with the right MFG as seed region
| Anatomical region | Cluster<br>size | MNI coordinates (mm) |  |  | *T*-value |
| --- | --- | --- | --- | --- | --- |
|  |  | *x* | *y* | *z* |  |
| Left cerebellum | 176 | -20 | -44 | -32 | 5.26 |
| Left cerebellum |  | -26 | -34 | -32 | 5.07 |
| Brainstem |  | -14 | -34 | -32 | 3.50 |

| Anatomical region | Cluster<br>size | MNI coordinates (mm) |  |  | *T*-value |
| --- | --- | --- | --- | --- | --- |

|  |  | **x** | **y** | **z** |  |
| --- | --- | --- | --- | --- | --- |
| Left cuneus | 307 | -14 | -82 | 16 | 4.74 |
| Left superior occipital<br>gyrus |  | -18 | -98 | 16 | 4.44 |
| Left superior occipital<br>gyrus |  | -14 | -92 | 12 | 3.99 |

| **Anatomical region** | **Cluster** | **MNI coordinates (mm)** |  |  | **T-value** |
| --- | --- | --- | --- | --- | --- |
|  | **size** |  |  |  |  |
|  |  | **x** | **y** | **z** |  |
| White matter | 153 | 18 | -22 | 36 | 4.94 |
| Middle cingulate<br>gyrus |  | 14 | -16 | 40 | 4.08 |
| Posterior cingulate<br>gyrus |  | 14 | -26 | 40 | 3.63 |

| Anatomical region | Cluster size | MNI coordinates (mm) |  |  | *T*-value |
| --- | --- | --- | --- | --- | --- |
|  |  | *x* | *y* | *z* |  |
| Right calcarine sulcus | 155 | 10 | -70 | 6 | 5.24 |
| Right calcarine sulcus |  | 10 | -80 | 10 | 4.17 |
| Right lingual gyrus |  | 2 | -70 | 4 | 3.97 |

Uncorrected $p < 0.001$ at peak level, false discovery rate-corrected $p < 0.05$ at the cluster level. MNI,
Montreal Neurological Institute

Regarding the β_art_, there were significant FCs between the right dorsal agranular insula and the middle/posterior cingulate cortex (MCC/PCC), and between the right ventral agranular insula and the right visual cortex (Figure 11C–D, Table 6). In contrast, we found no brain region showing significant FC with the regions adjacent to the anterior insula.

## Discussion

In this study, we examined the neural basis of anxiety and anxiety-related autonomic responses during a daily driving situation. We hypothesized that the brain network related to the elicitation of anxiety and corresponding autonomic responses would be observed in the anterior insula and subcortical regions. As predicted, during the period in which the participants anticipated a car crash, we observed cortico-subcortical activations in the anterior insula, PAG, thalamus, and BNST. The amygdala activity observed in response to a crash (but not detected in anticipation of a crash) could extend our understanding of the role of the amygdala in the elicitation of anxiety. Moreover, we observed deactivation in the posterior insula and SII, which could reflect an allostatic response against the upcoming crash. This could offer new insights for the safe-signal-related brain network that has been unclear from previous fear-conditioning studies. We also observed significant pupil dilation, although the increase in β_art_ related to the anticipation of a crash was not statistically significant. The FC analyses using these autonomic responses revealed that β_art_-related FC was found between the right dorsal anterior insula and cingulate cortex, as well as between the right ventral anterior insula and visual cortex, and that pupil-related FC was found between the right MFG and the cerebellum/brainstem, as well as between the right rostral IFG and the visual cortex. Our results could extend the knowledge regarding the neural basis of anxiety and anxiety-related physiological responses.

### Brain regions involved in anxiety elicitation

In the attention condition, the brain regions including the anterior insula, PAG, thalamus, and BNST were more active than those in the safe condition. The anterior insula is reportedly involved in the integration and prediction of interoceptive information (Craig 2009) and in the allostatic process based on prediction (Barrett et al. 2017). The peak voxel of the anterior insula cluster is located at the dorsal part of the anterior insula and extends to the operculum and triangular parts of the IFG. These areas are consistent with those related to sustained fear reported by Somerville et al. (2013). The dorsal anterior insula has been suggested to be involved in cognitive and attentional functions rather than in emotional function (e.g., Chang et al. 2013). This could suggest that the anterior insular activity during the attention condition could reflect only the attentional process; however, based on the results of this study, such a suggestion can be ruled out. The results of anxiety ratings indicated that participants felt more anxious during the attention condition than during the safe condition. In contrast, RTs showed no significant difference between conditions (see Figure 3), suggesting that attentional resources allocated to the task were similar among the conditions. Furthermore, we found no significant difference in activation in the dorsal attentional network, such as the dorsolateral prefrontal cortex, frontal eye field, and posterior parietal cortex, between the attention and safe conditions. Thus, our results suggest that the brain regions including the anterior insula were related to anxiety, that is, negative emotion caused by prediction of an aversive event rather than just attention.

The PAG receives projections from the prefrontal cortex, insula, and amygdala (Mantyh 1982). The PAG is suggested to play a role in homeostatic defense by integrating afferent information from the periphery and information from higher centers (Linnman et al. 2012). For instance, the PAG is known to be involved in pain modulation. It has been reported that when pre-stimulus FC between the anterior insula and PAG is stronger, participants perceive a pain stimulus as less painful (Ploner et al. 2010).

The thalamus is also reportedly involved in the modulation of ascending nociceptive information (Tang et al. 2009). Moreover, it is known that the thalamus relays nociceptive information to the insula, anterior cingulate cortex, and somatosensory cortices (Kummer et al. 2020). In the current study, we observed thalamic activation mainly in the dorsomedial region. Considering that the dorsomedial thalamus is a part of the medial spinothalamocortical pathway involved in nociceptive-specific responses (e.g., Gingold et al. 1991), the thalamic activity observed in this study presumably reflects the anticipatory modulation of responses to unpleasant stimuli accompanied by collision.

The BNST is known to be involved in emotion, threat, and autonomic processing (Davis et al. 2010; Crestani et al. 2013). While animal studies have reported the structural connection between the insula and BNST (e.g., Centanni et al. 2019), the BNST-insula connection in human brains has also been revealed in recent MRI studies (e.g., Avery et al. 2014; Flook et al. 2020). This connection is suggested to be involved in the translation of emotional states into behavioral responses, including “fight or flight” (Flook et al. 2020). Based on these previous studies, activation in the BNST and anterior insula detected in the attention condition might be involved in the elicitation of autonomic and behavioral responses accompanied with anticipation of a crash. Thus, our results suggest that a cortico-subcortical network comprising the anterior insula, PAG, BNST, and thalamus is involved in the prediction of negative events and elicitation of anxiety-related responses based on interoceptive information in the attention condition.

In contrast, the SII/posterior insula and precuneus were more activated in the safe condition than in the attention condition. In the attention condition, where a crash was predicted, the activation in these regions decreased as the timing of the crash approached (see Figure 8C). Therefore, this deactivation could reflect the allostatic process preparing for the predicted response to a possible upcoming aversive event. The SII/posterior insula has been reported to be involved in processing aversive stimuli (e.g., Apkarian et al. 2005). In particular, the SII has been suggested to represent somatosensory prediction error (Blakemore et al. 1998). In the study by Blakemore et al., participants underwent two experimental conditions: the experimenter tickled the participants’ palms, and the participants tickled themselves. In the condition of self-tickling, the SII was more deactivated than in the condition in which the experimenter tickled the participants, suggesting that this deactivation reflects the process of canceling out the sensory consequences, that is, a tickly feeling, generated by prediction. In our study, crash-related activation overlapped in the SII/posterior insula cluster, as observed in the contrast of safe > attention. This suggests that SII/posterior insula activation could reflect a part of the aversive sensation elicited by the crash. Considering these observations, the deactivation in the SII/posterior insula observed in the attention condition could be interpreted as an allostatic attenuation triggered by the prediction of sensory input followed by an upcoming aversive event.

The precuneus is part of the DMN. It has been proposed that the DMN is switched to the central executive network according to the task demand by the salience network (Menon and Uddin 2010). Therefore, deactivation in the precuneus could reflect inhibition of the DMN to prepare and pay attention to the upcoming event in the attention condition.

### Brain activity related to subjective anxiety

The results of the parametric modulation analyses showed that subjective anxiety was significantly correlated with activation in the thalamus and PAG, and not with the anterior insula. This could be explained by the relationship between anterior insula-PAG pre-stimulus FC and subjective pain sensation (Ploner et al. 2010). Considering that the anterior insula-PAG connection during the anticipation period determines subjective anxiety, when anticipatory modulation of PAG by the anterior insula was insufficient, the participants could have felt more anxious. In contrast, regardless of subjective anxiety, the anterior insula was active when an upcoming crash could be predicted, which could contribute to the result that the anterior insula was less related to subjective anxiety.

While many previous studies have suggested that the amygdala is involved in anxious emotion, we did not find significant amygdala activity during the anticipation period in the contrast of attention > safe or in the parametric modulation of subjective anxiety. However, the amygdala was significantly active in response to a crash relative to the baseline (Figure 10B)^1^. This is consistent with the study of Somerville et al. (2013), who reported that the amygdala responds to transient fear rather than sustained anxious emotion (anxiety). Thus, in the current study, the absence of amygdala activity during the anticipation phase is reasonable.

### Brain regions involved in anxiety-related autonomic responses

The autonomic response-related FC analyses revealed that the right anterior insula and its adjacent regions showed significant FC with the visual cortex, cerebellum, brainstem, and MCC/PCC.

Related to β_art_, the right ventral and dorsal anterior insula showed FC with the visual cortex and MCC/PCC, respectively. The anterior insula controls the autonomic nervous system by regulating the subcortical brain areas, such as the PAG (Thayer and Lane 2009; Critchley and Harrison 2013). Previous research has demonstrated that autonomic control of cardiac activity is lateralized and mediated by the right insular cortex (Colivicchi et al. 2004; Craig 2009). For instance, evidence from stroke studies has suggested that the right insula plays a major role in cardiac autonomic control (e.g., Colivicchi et al. 2004). Craig (2009) proposed a hypothesis that the right and left insular cortices are involved in the control of the sympathetic and parasympathetic nervous systems, respectively. Based on these findings, the observation of β_art_-related FC for the right anterior insula as seed region is reasonable. The MCC/PCC cluster showing FC with the right dorsal anterior insula was located at the posterior part of the MCC and PCC. Particularly, the PCC is suggested to be involved in controlling the balance between an internal and external attentional focus in collaboration with the right anterior insula (Leech and Sharp 2014). The ventral anterior insula, which is involved in affective processing, showed FC with the visual cortex. This might reflect the allostatic processing of upcoming visual information based on anxiety elicitation. These observations suggest that the allostatic affective processing was associated with the β_art_ change. However, Tsuji et al. (2021) demonstrated that the β_art_ during pain stimulation was correlated with brain activity in the salience network, including the dACC but not the anterior insula. This inconsistency could suggest that the anterior insula is related to the β_art_ attributed to anticipation, whereas the dACC is related to that caused by stimulation.

Regarding the pupil-related FC, the right ventral MFG and the rostral IFG showed FC with the cerebellum/brainstem and the visual cortex, respectively. The brain regions including the anterior insula/IFG and MFG have also been reported to be related to pupil change during reward anticipation (Schneider et al. 2018). The ventral MFG and rostral IFG, which are adjacent to the inferior frontal sulcus, are a part of the ventral attention network (Corbetta et al. 2008). The ventral attention network is known to be a part of the salience network and has been suggested to be a “circuit breaker” that reorients focused attention by detecting salient stimuli such as oddballs. The ventral MFG showed an FC with the cluster including the pons located near (however, not including) the locus coeruleus (LC), which is involved in pupil control (Schneider et al. 2016). Furthermore, FC between the IFG and visual cortex was observed. This FC could reflect allostatic attentional control to visual information. Considering that no pupil-related FC was observed with the anterior insula as seed, the pupil-related FC in the current study could reflect a relatively cognitive aspect of allostatic processing, such as attentional control to visual information accompanied by an upcoming collision.

Thus, our results suggest that the anterior insula and its adjacent regions, in collaboration with other regions, could differentially realize an allostatic attention control and adaptive physiological response to environmental information.

However, we did not observe FC with the brain regions involved in general direct control of the pupil and artery, such as the LC for pupil dilation and the hypothalamus for the artery. One possible interpretation of this result is that the regions observed in this study could be involved in task-evoked modulation of the pupil and artery as part of the allostatic process. During the resting state and the continuous attentive task, for instance, the pupil diameter is known to be correlated with BOLD signals in the LC (Murphy et al. 2014). However, certain studies have reported that LC activation is not related to task-evoked pupil dilation (Schneider et al. 2018). Nevertheless, the relationship between the task-evoked pupil- and artery-related regions and those involved in the general control of the pupil and artery should be further examined in future research.

Regarding pupil dilation, we found a robust difference between the attention/crash and safe conditions. This result suggests the possibility that the measurement of pupil dilation is useful in detecting anxiety. However, we must note that the correlation between the pupil dilation and subjective ratings of anxiety were not significant and that we also observed a weak (but not significant) correlation between pupil dilation and subjective arousal. Therefore, future studies controlling the arousal level should assess whether pupil dilation merely reflects the arousal.

In contrast, the increase of β_art_ in the attention/crash conditions relative to the safe condition was not significant. This result could have been caused by the difference in the temporal properties of pupil dilation and β_art_. Arousal-related pupil dilation reportedly appears at nearly 200 ms (Sirois et al. 2014). In the current study, owing to the fast latency of pupil dilation, the fluctuation in the participants’ subjective emotional state could be reflected more accurately than β_art_. In contrast to pupil dilation, β_art_ is a relatively newly proposed index. Information on the temporal properties and mechanism of β_art_ should be investigated in future studies.

### Limitations and future directions

In the current study, we demonstrated that the network including the anterior insula plays a pivotal role for eliciting anxiety and for accompanying autonomic responses in a daily situation. However, our results do not elucidate clinical conditions, such as anxiety disorder. For instance, we found no significant correlation between the trait and state anxiety and the subjective ratings. Furthermore, there was no significant brain activation related to the individual difference in the trait anxiety nor the state anxiety before and after the experiment, as assessed with the STAI. This could be attributed to the fact that we only recruited healthy participants with a relatively narrow range of scores. To apply our findings to the understanding of anxiety disorder, one should compare brain activity and physiological responses between patients and normal controls. One of the aims of future studies should be to assess whether anxiety-prone individuals exhibit an increase in the pupil area and β_art_, which will deepen our understanding of the mechanism of anxiety. In addition, we should note an imbalance in the male-female ratio in this study. Regarding gender differences in emotion, there has been a debate in the field of emotion studies. A review from Kret and De Gelder (2012) pointed out that men tend to show strong effects of threatening stimuli, such as enhanced physiological arousal and brain activity. The gender difference of the anxiety-related brain and physiological responses should be examined in future studies.

There is also a limitation to the localization of the small subcortical regions, such as the BNST and PAG. Although we confirmed that the activated cluster included the BNST and PAG using anatomical atlases, more precise localization is necessary for a better understanding of the roles of the nuclei in anxiety-related responses in future. Such may be achieved with specific scan parameters to target the small nuclei or MRI with a higher magnetic field (e.g., 7-T MRI).

Our results also showed that the increase in pupil area and peripheral arterial stiffness were commonly related to the network including the right anterior insula and its adjacent regions in a daily situation of driving a vehicle. This suggests that these are potential indices to detect the driver’s anxiety during driving, underpinned by neuroscientific evidence. If our results are applied to daily driving situations, however, there are some limitations. First, the task was a simple fear-conditioning task, which did not contain actual control of a handle, accelerator, or brake, as in an actual driving situation. Moreover, we did not perform a comparison to a “non-driving” condition. Second, the experimental environment in the MRI scanner differs much from an actual driving situation. For instance, the participants did not hold a steering wheel or place their foot on a brake pedal. Furthermore, they were lying in a supine position in the MRI scanner. For industrial application, an experiment should be conducted with a realistic steering device (e.g., Okamoto et al. 2020) that can be used in an MRI scanner, enabling a more realistic driving situation, and a non-driving condition should be included for comparison. It is also necessary to perform brain imaging and physiological recordings using wearable devices (e.g., Protzak and Gramann 2018) while driving an actual vehicle.

## Conclusions

Our results suggest that cortico-subcortical regions, including the right anterior insula, PAG, thalamus, and BNST, play a core role in anticipatory responses to upcoming threats in daily driving situations. We also observed that multiple autonomic responses, such as pupil dilation and β_art_, were evident when an upcoming threat was predicted, and the right anterior insula and its adjacent regions play a key role in eliciting autonomic responses. These results suggest that pupil dilation and β_art_ reflect anxiety-related salience network activity, and that they are potential candidate indices for detecting task-evoked anxious emotion based on neuroscientific evidence.

## Acknowledgments

We are grateful to Dr. Norihiro Sadato, Dr. Alan Fermin, Dr. Kentaro Ono, Dr. Ayumu Matani, Dr. Toshio Tsuji, and Dr. Maro G. Machizawa for helpful discussions concerning this study, and Ms. Noriko Miura and Ms. Tamami Tomita for assistance with the implementation of the experiment. We would like to thank Editage (www.editage.com) for English language editing.

## Footnotes

1 We must note that the crash condition was quite different from other conditions, as indicated in the behavioral data. For example, the RTs to the word “STOP” in the crash condition were faster than in other conditions. This could be attributed to the auditory-visual stimuli accompanied with a crash that captured the participants’ attention, thereby enhancing their responses to the word “STOP.” Furthermore, the subjective ratings for the crash condition were significantly different from those for other conditions, indicating that the subjective ratings performed after each trial were affected by the crash, as predicted.

