## Supplementary Materials for "Neural basis for anxiety and anxiety-related physiological responses during a driving situation: An fMRI study"

### 1 Supplementary Figure

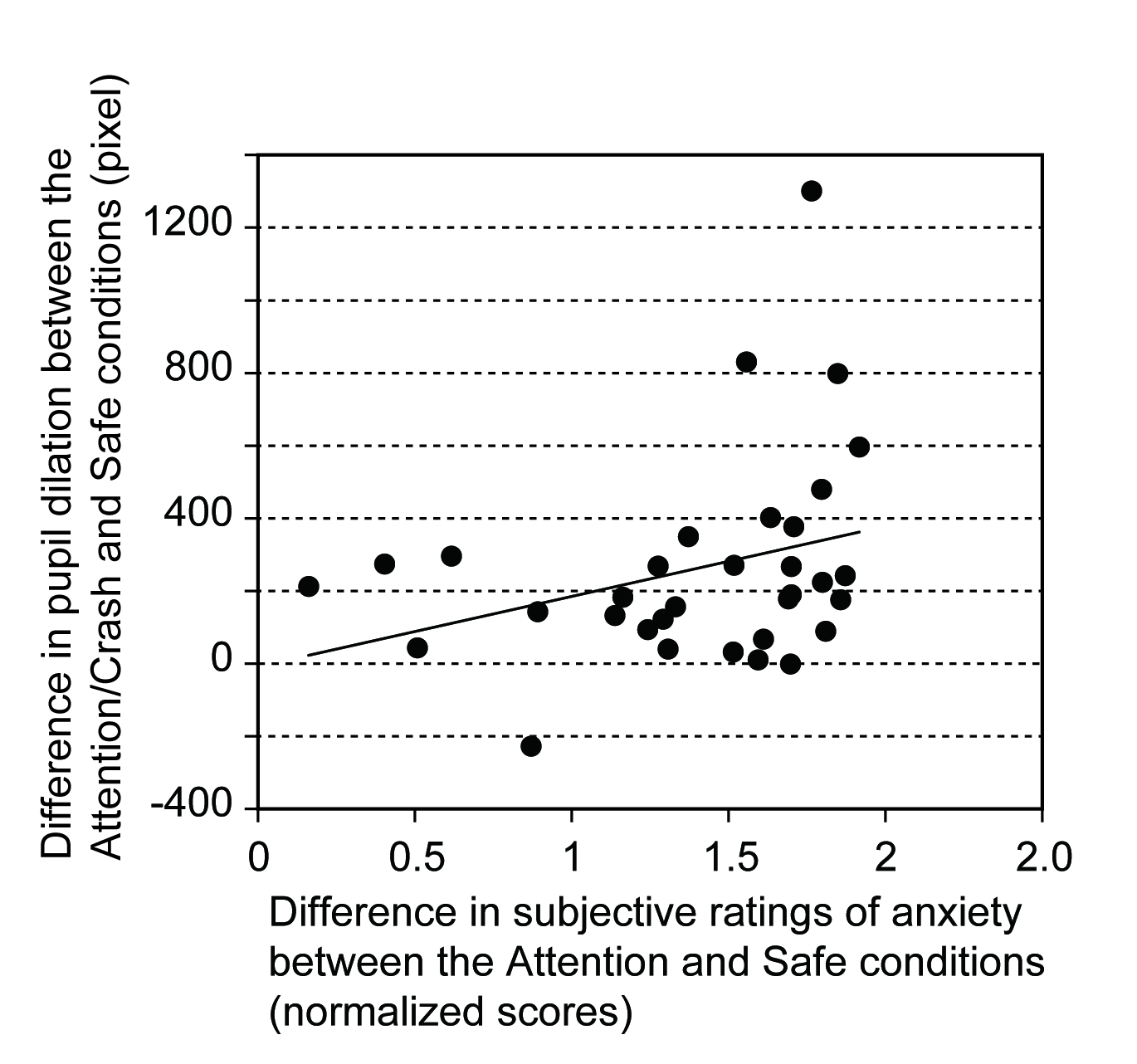

**Supplementary Fig. S1.** Pupil dilation and anxiety ratings. The correlation analysis in pupil size between the “attention/crash” and “safe” conditions, and the subjective ratings of anxiety between the “attention” and “safe” conditions. The line represents the result of a linear regression.
